# CRISpy-pop: a web tool for designing CRISPR/Cas9-driven genetic modifications in diverse populations

**DOI:** 10.1101/2020.06.19.162099

**Authors:** Hayley R. Stoneman, Russell L. Wrobel, Michael Place, Michael Graham, David J. Krause, Matteo De Chiara, Gianni Liti, Joseph Schacherer, Robert Landick, Audrey P. Gasch, Trey K. Sato, Chris Todd Hittinger

## Abstract

CRISPR/Cas9 is a powerful tool for editing genomes, but design decisions are generally made with respect to a single reference genome. With population genomic data becoming available for an increasing number of model organisms, researchers are interested in manipulating multiple strains and lines. CRISpy-pop is a web application that generates and filters guide RNA sequences for CRISPR/Cas9 genome editing for diverse yeast and bacterial strains. The current implementation designs and predicts the activity of guide RNAs against more than 1000 *Saccharomyces cerevisiae* genomes, including 167 strains frequently used in bioenergy research. *Zymomonas mobilis*, an increasingly popular bacterial bioenergy research model, is also supported. CRISpy-pop is available as a web application (https://CRISpy-pop.glbrc.org/) with an intuitive graphical user interface. CRISpy-pop also cross-references the human genome to allow users to avoid the selection of sgRNAs with potential biosafety concerns. Additionally, CRISpy-pop predicts the strain coverage of each guide RNA within the supported strain sets, which aids in functional population genetic studies. Finally, we validate how CRISpy-pop can accurately predict the activity of guide RNAs across strains using population genomic data.

## Introduction

CRISPR/Cas9 has become a widely used genome-editing tool due to its accuracy, precision, and flexibility (Ceasar *et al*. 2016). A primary step in designing a CRISPR/Cas9 experiment is the selection of the single guide RNA (sgRNA) target site, which usually occurs twenty nucleotides upstream of a protospacer adjacent motif (PAM) site. This sgRNA is bound by the Cas9 enzyme and used to direct Cas9 to the complementary location within the genome. Once bound to DNA, the Cas9 endonuclease cuts the DNA, leaving a double-strand break in the chromosome. This break can then be repaired using nonhomologous end-joining or homology-directed repair. Modified versions of Cas9 can nick a single DNA strain or bind to DNA sequence motifs without cleaving either DNA strand, while other Cas proteins have different sequence requirements (Makarova and Koonin 2015). Due to the ease of manipulation of sgRNA targets by Cas9, it has been widely used for genome editing, including in yeasts and bacteria used in bioenergy research (Wang *et al*. 2016, Huang *et al*. 2016, and Dong *et al*. 2016, Higgins *et al*. 2018, Kuang *et al*. 2018). For *Zymomona mobilis*, there have been more published successes using Cas12a, which uses a different PAM site than Cas9 (Shen *et al*. 2019).

When designing sgRNAs, two main considerations must be made: efficiency and specificity. One tool that addresses the prediction of sgRNA efficiency is sgRNA Scorer 2.0 (Chari *et al*. 2017), which generated a model across multiple Cas9 orthologs to predict activity of sgRNAs from their sequence composition. A tool that addresses specificity of sgRNAs is Cas-OFFinder (Bae *et al*. 2014), which is a fast algorithm that searches specific genomes for potential off-target sites. Cui *et al*. 2018 reviewed a panel of twenty representative sgRNA design tools, which vary in their genome specificity, nuclease(s) supported, user input, and methods (or lack thereof) for on-target prediction and off-target scoring. Of the reviewed tools, eight supported the *Saccharomyces cerevisiae* reference genome and allowed for a variety of PAM sites. Of those eight, six provided both on-target prediction and off-target scoring, but none combined the use of sgRNA Scorer 2.0 and Cas-OFFinder into a single tool.

Recent advances in high-throughput sequencing have enabled the collection of population genomic data for an increasing number of organisms. Many studies have sequenced whole genomes of traditional and emerging model organisms, including large populations. For example, the 1000 Genomes Project (Auton *et al*. 2015) sequenced human genomes, the 1001 Genomes Project (Alonso-Blanco *et al*. 2016) sequenced *Arabidopsis thaliana* genomes, and the 1002 Genomes Project (Peter *et al*. 2018) sequenced *S. cerevisiae* genomes. With the increasing availability of population genomic data and the need to determine the functions of polymorphisms, there is a growing need to accommodate variation within species when designing CRISPR/Cas9-driven genetic modifications. Recently, the SNP-CRISPR tool was developed to address genomic variation by targeting single nucleotide polymorphisms (SNPs) (Chen *et al*. 2020). SNP-CRISPR supports several genetic model organisms, including humans, mouse, and *Drosophila melanogaster*, but it is limited to the variants included in user-supplied files, which creates a barrier for less computationally proficient users. To our knowledge, neither this tool, nor any other existing tool, supports *S. cerevisiae* population genomic datasets.

Here we developed and describe CRISpy-pop as a python-based (Van Rossum and Drake 2009) web application for the design of <u>CRIS</u>PR/Cas9 sgRNAs for genetic modifications on <u>pop</u>ulations of strains. CRISpy-pop incorporates popular diverse strain sets of *S. cerevisiae* from recent population genomic studies (Peter *et al*. 2018; Sardi *et al*. 2018) and uses the existing tools sgRNA Scorer 2.0 and Cas-OFFInder to assess the strain coverages of sgRNAs, predict their activities, and determine their off-target potentials. As a proof of principle, here we use CRISpy-pop to design *ade2* knockout mutants and accurately predict which strains can be targeted by which sgRNAs. CRISpy-pop fills a needed niche in functional and population genomic research.

## Materials and Methods

### CRISpy-pop pipeline

The CRISpy-pop bioinformatic pipeline supports three modes of operation: targeting a gene, offsite target search, and targeting a custom sequence (**Figure 1**). We made use of open-source bioinformatic tools to generate sgRNA designs. The resulting sgRNA sequences are then scored and ranked based on predicted efficiency of the sgRNAs. Offsite target interactions are reported for each sgRNA across two complete strain sets of *S. cerevisiae*. These results are displayed to the user in a convenient and intuitive graphical user interface (GUI). The user can sort, search, save, and export the results in a more efficient way than would be possible using command line tools alone. CRISpy-pop also contains a genome viewer for visualization of each sgRNA within the target gene, facilitating design choices for the desired genome edits. CRISpy-pop supports a 167-strain set of *S. cerevisiae*, including 165 recently published genomes (Sardi *et al*. 2018), the S288C reference genome (Engel *et al*. 2014), and the GLBRCY22-3 bioenergy chassis (McIlwain *et al*. 2016); as well as a 1011-strain set of *S. cerevisiae* from the 1002 Yeast Genomes Project (Peter *et al*. 2018). CRISpy-pop’s population genetic tool reports the numbers and identities of strains with perfect matches to each sgRNA. CRISpy-pop also contains a biosafety feature, which performs a local BLAST search (Altschul *et al*. 1990) of the human genome for perfect matches to each sgRNA sequence.

**Figure 1.**
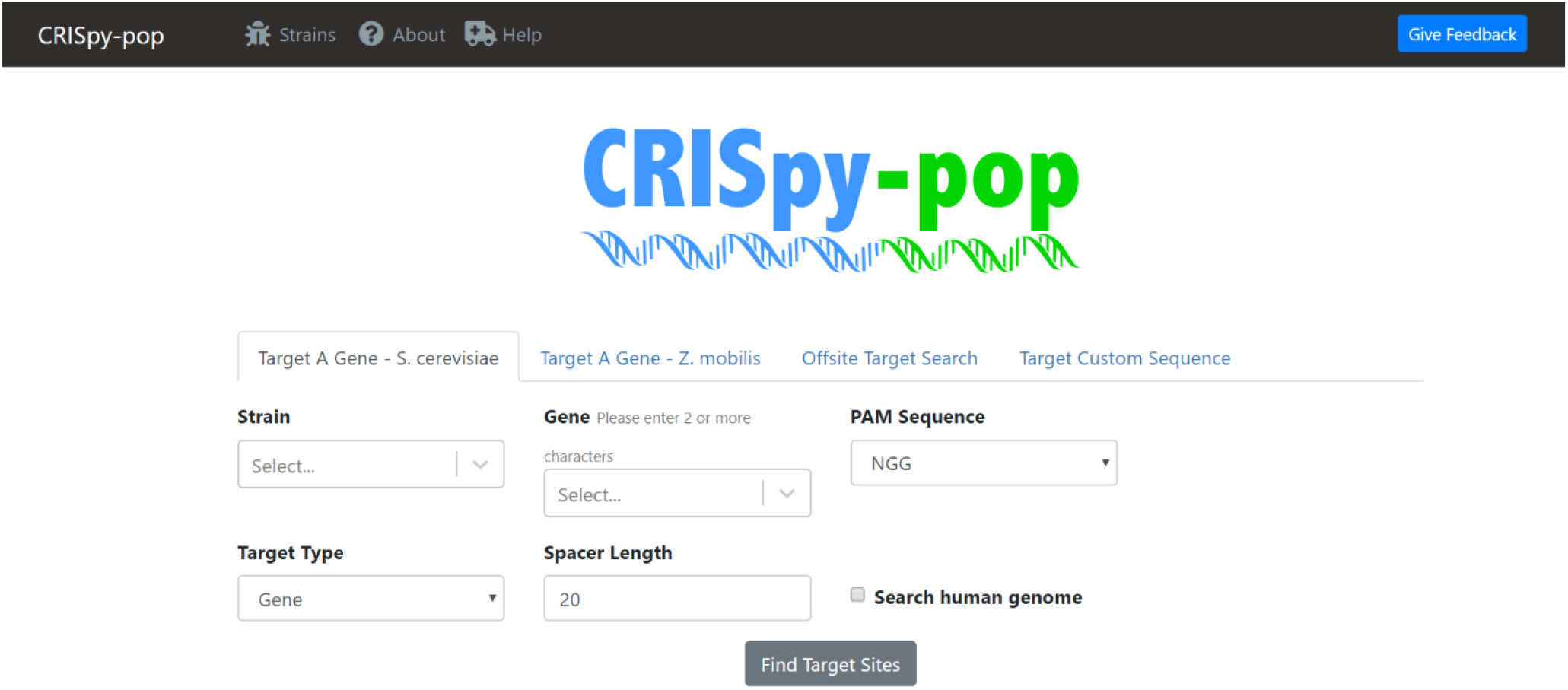
Screenshot of the CRISpy-pop homepage (https://CRISpy-pop.glbrc.org/). There are options to search a gene in *S. cerevisiae* and *Z. mobilis*, as well as an offsite and custom target search. There are options to select specific strains, the desired PAM site, and the sgRNA length. Users may select the following PAM sites: NGG, NNGRRT, TTTV, NNNNGATT, or NNAGAAW. Additionally, there is an option to search the human genome for perfect matches. CRISpy-pop features a user-friendly, web-based GUI.

### Targeting a gene in a specific strain

A mode was designed within CRISpy-pop to give the user the ability to target a gene by name in a specific strain. When the user selects this option, a streamlined search is performed to generate sgRNA sequences as follows. Gene coordinates are extracted from the appropriate GFF file. Using the reference genome FASTA, the gene sequence is extracted using samtools (Li *et al*. 2009). CRISpy-pop uses VCF files from its internal set of strains. These VCF files each contain variant calls for each strain relative to the S288C reference genome. These variants are then used to make substitutions in the S288C genome sequence to produce sequence files for each strain. This sequence is used as input to sgRNA Scorer 2.0 (Chari *et al*. 2017) in FASTA format with the appropriate PAM sequence and orientation (5’ or 3’) and the desired sequence length, which outputs a list of sgRNA sequences and their predicted activity scores. Cas-OFFinder (Bae *et al*. 2014) is then used to query all strains for offsite interactions, allowing zero mismatches. The results are output in a user-friendly, graphical format. The results include a genome viewer, which shows the relative position of each sgRNA for the gene. In a table format, for each sgRNA, CRISpy-pop reports the sgRNA sequence, PAM site, activity score, GC%, chromosome, position, strand, position in the gene, mismatches, off-site matches, human genome hits, and strain coverage. Specific information can be obtained for any individual sgRNA by clicking on the desired entry in the table. For each sgRNA, an individual result report can be viewed, containing identities of strains predicted to be targeted, the alignment with the target, and the sgRNA details and statistics.

### Offsite target search

For the offsite target search mode, a user provides a previously designed sgRNA sequence and selects the reference genome to be searched. Upon each search, CRISpy-pop employs Cas-OFFinder to provide a list of the specified reference genome’s offsite targets for the user specified sgRNA. If no offsite targets exist, CRISpy-pop outputs that none were found.

### Target a custom sequence

This mode allows the user to target a custom sequence, such as a gene that they may have previously engineered into a strain. When this feature is used, a custom DNA sequence is entered by the user. Once this sequence is entered, CRISpy-pop uses sgRNA Scorer 2.0 to find and score all potential sgRNAs within that sequence. Optionally, several supported reference genomes can be searched for offsite target matches, again using Cas-OFFinder. Currently supported genomes include *S. cerevisiae* S288C, *S. cerevisiae* GLBRCY22-3, *Saccharomyces paradoxus, Kluyveromyces lactis*, and *Zymomonas mobilis* ZM4. This tool outputs the same results as the gene target search.

### Human hits search

CRISpy-pop performs a BLASTn database search of the human genome version hg38 (Schneider *et al*. 2017) for exact matches to each sgRNA as the query. If any perfect matches to the sgRNA are found, the output reports “Yes” under human hits.

### Strain coverage function

This function uses the population genomic data described above to determine which strains are predicted to be targeted by each sgRNA. The strain coverage function searches the selected strain set for perfect matches to the sgRNA sequence and reports the number and identities of the strains covered.

### ADE2 sgRNA selection

To validate CRISpy-pop’s functionality, we used it to find sgRNAs to target the gene *ADE2* using a spacer length 20 and a PAM site of NGG. The 167-strain set (165 isolates, S288C and GLBRCY22-3) was searched for strain coverage. Two sgRNAs were selected with high strain coverage, and two were selected with low strain coverage, the latter of which were selected to have the exact same strain identities covered. The sgRNAs were chosen to balance the need for high activity scores, target more 5’ positions within the gene, and have no offsite matches.

**Table 1.**
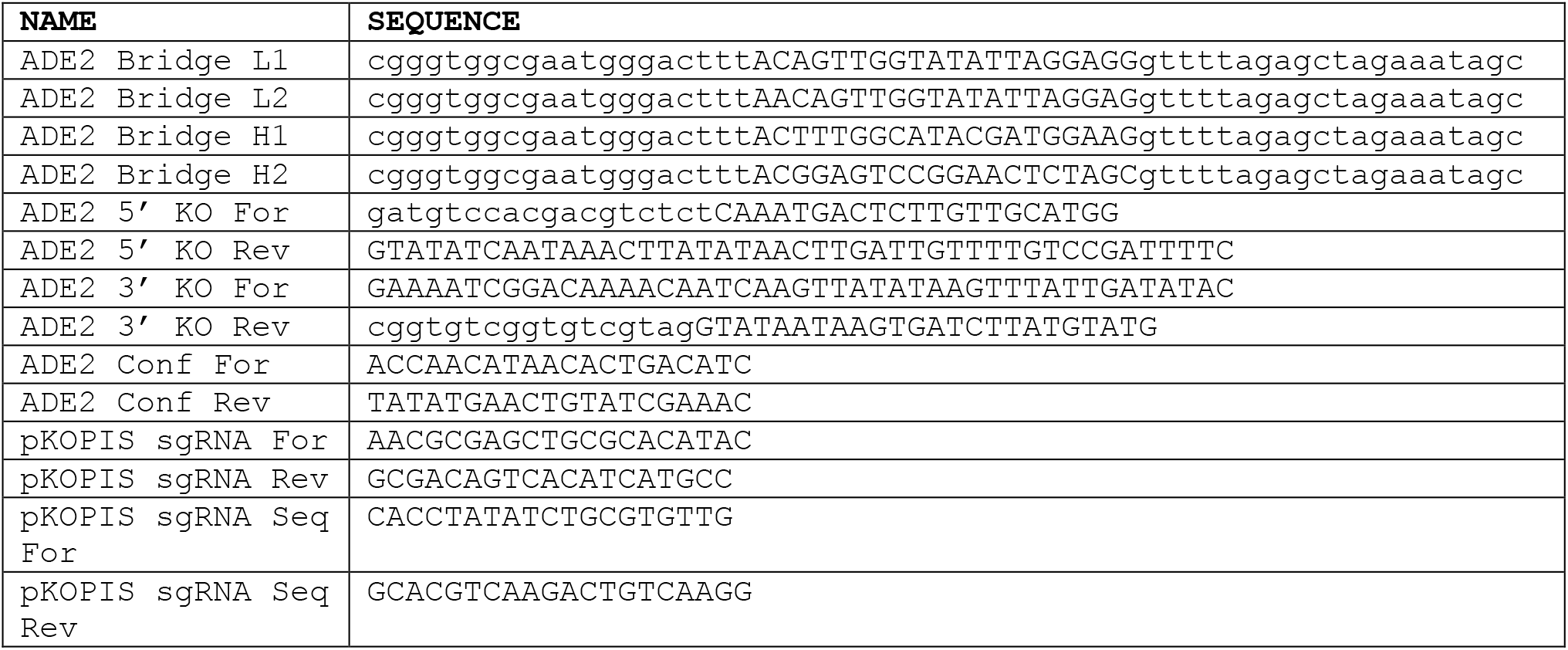
Table of the oligonucleotides used. These include the bridge primers for adding the sgRNA sequences to the pKOPIS + sgRNA plasmid, the primers for PCR SOEing to clone the donor DNA, and the primers for PCR and Sanger sequencing.

### Plasmid and donor DNA synthesis

An empty sgRNA expression cassette, which contained the *SNR52* promoter, HDV ribozyme linked to a cloning site for sgRNA construct, and the *SNR52-1* terminator (Kuang *et al*. 2018), was first cloned into the pKOPIS plasmid (Kuang *et al*. 2018) using the NEBuilder HiFi DNA Assembly Master Mix (NEB #E2621) (Hsieh 2018). pKOPIS contains a *kanMX* selectable marker and encodes a Cas9 protein driven by the constitutive *RNR2* promoter. This empty pKOPIS + sgRNA plasmid (pHRW68) was linearized using a restriction enzyme digest with *Not*I.

Four different 60-nucleotide (nt), single-stranded bridging primers were designed, each containing one of the selected sgRNA sequences flanked by 20-nt homology regions with the pKOPIS plasmid (**Table 1**). The NEBuilder HiFi DNA Assembly Master Mix (Hseih 2018) was then used to clone the sgRNA sequences into the pKOPIS+ sgRNA plasmid, using the linearized plasmid and the bridge primers. This mixture was then used to transform *Escherichia coli* cells. The plasmids with the inserted sgRNAs were each isolated using the ZR Plasmid Miniprep Classic kit (Zymo Research). We confirmed correct sgRNA sequence insertion by performing BigDye™ (Applied Biosystems) Sanger-sequencing reactions with the pKOPIS sgRNA Seq primers.

The donor DNA was constructed using PCR splicing by overlap extension (SOEing) (Horton *et al*. 2013). All but the first 100 and last 100 base pairs of the gene were designed to be deleted from *ADE2*. A 40-nt primer was designed to amplify the 5’ forward portion of the gene and the homology region. A 40-nt primer was designed to amplify the 5’ reverse portion of the gene with 20-nt from the first 100 base pairs (bp) of the gene and 20-nt from the last 100 bp. The complement of this primer was then designed to amplify the 3’ forward portion of the gene. Finally, a 40-nt primer was designed for the 3’ reverse portion of the gene, which contained the last 20-nt of the gene and the homology region. Additionally, ADE2 Conf FOR and ADE2 Conf REV primers (**Table 1**) were used to confirm deletion of *ADE2* by PCR and sequencing.

The 3’ and 5’ sections of the donor DNA were first amplified individually using gradient PCR with annealing temperatures from 50°C – 70°C and Phusion^®^ High-Fidelity DNA Polymerase (New England Biolabs). The two individual sections were then joined into the complete donor DNA fragment using the same gradient PCR protocol. The final product was purified using the Qiagen QIAquick PCR Purification Kit according to the manufacturer’ s directions. (Qiagen).

### Transformation and knockout screening

All transformations were performed with pKOPIS plasmids containing each of the four sgRNAs or the empty vector as a negative control using the standard lithium acetate protocol optimized for *S. cerevisiae* (Gietz *et al*. 1995). In each reaction, 0.75 μg of sgRNA and 2 μg donor DNA were used. The transformations were grown in liquid YPD for three to five hours at 30° on a tissue culture rotator. They were then plated on three YPD + G418 (200μg/L) plates, with 100μl, 200μL, and 300μL of transformation per plate. Successful transformants grew on YPD + G418 plates, while successful *ade2* knockouts also turned pink. The total number of colonies on each YPD + G418 plate was counted, as well as the number of pink colonies. The number of pink colonies was divided by the total number of successful transformants to calculate the efficiency of *ade2* deletion. Two pink colonies were chosen from each transformation of each strain, their *ADE2* genes were amplified by PCR, and their products were sequenced using Sanger sequencing to confirm that the knockouts had occurred using the donor DNA and homology-directed repair.

### Statistical analysis

For the two strains targeted by all four sgRNAs (K1 and S288C), we used Mstat (https://mcardle.oncology.wisc.edu/mstat/) to calculate Kendall’s Tau, performing a one-sided test for a correlation between the activity scores and the efficiencies.

### Data availability

CRISpy-pop is available online for non-commercial use at https://CRISpy-pop.glbrc.org/. The source code for the pipeline is available at: https://github.com/GLBRC/CRISpy-pop/. All new data generated is contained within this manuscript.

## Results and Discussion

### Population-level variation in sgRNA target sites

*S. cerevisiae* is a useful genetic model system and bioengineering chassis due to its well-studied genome and ease of genetic manipulation. With growing population genomic datasets, functional investigations with CRISPR/Cas9 tools can now be extended beyond traditional laboratory strains, but variation in sgRNA target sites can still limit portability. To explore variation within the 1011-strain set, we calculated the total number of genomes that could be targeted by each sgRNA (**Figure 2**). The total number of sgRNAs designed was 658,304. Only 73,495 of the sgRNAs have perfect matches in all 1011 genomes, while 584,809 sgRNAs are predicted to target some other fraction of the genomes. Thus, randomly picking a sgRNA designed against the S288C reference genome would be unlikely to target all strains of potential interest. CRISpy-pop allows users to sort and filter by the number of strains targeted in a given gene, which aids design decisions to maximize sgRNA portability and facilitates population-level studies.

**Figure 2.**
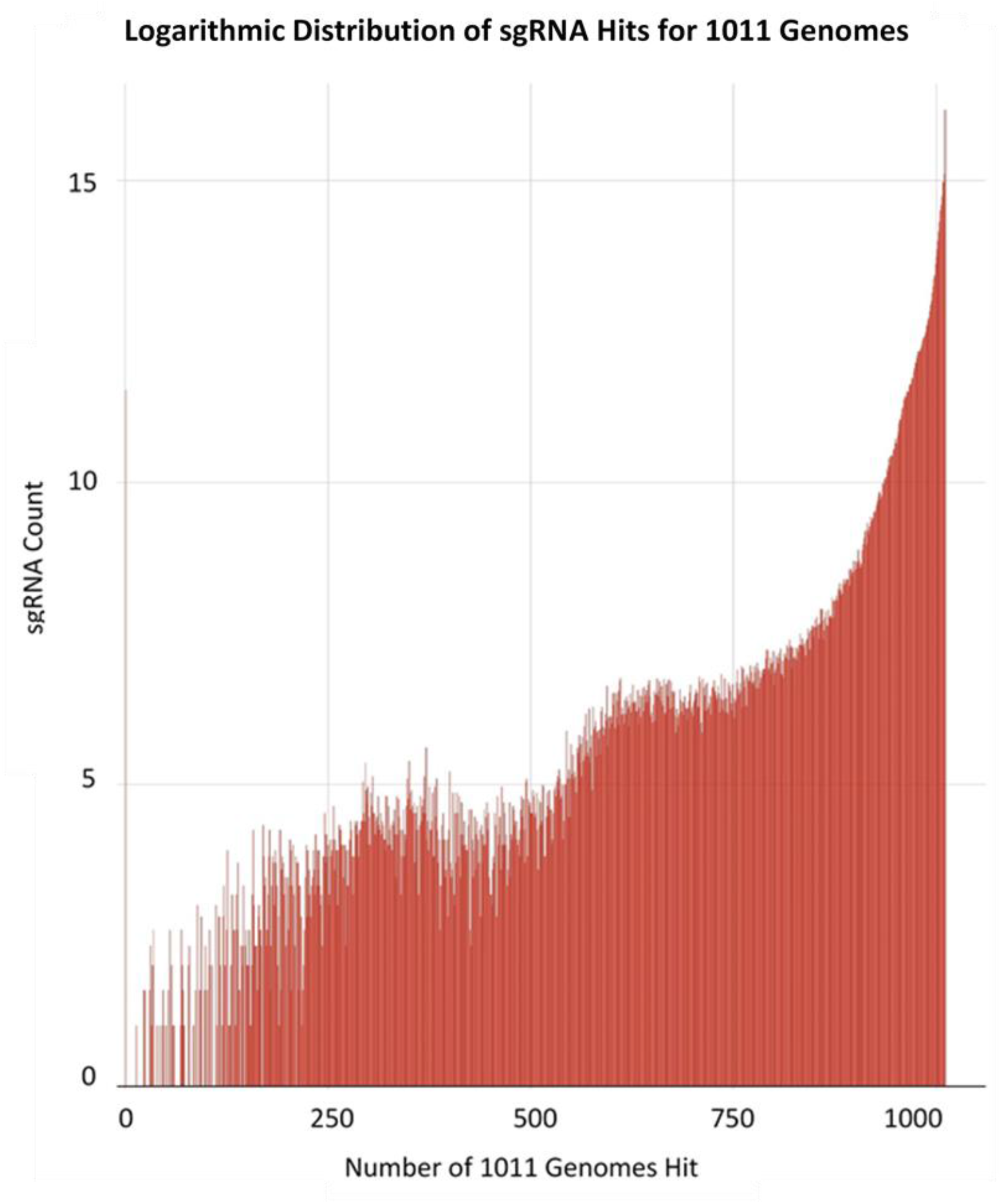
A histogram showing the number of sgRNAs, on a log_2_ scale, that target each number of the 1011-strain set. Only 73,495 sgRNAs are predicted to target all 1011 genomes, while the remaining 584,809 sgRNAs target only a fraction of the genomes.

### sgRNA selection using CRISpy-pop

To validate the strain coverage function of CRISpy-pop, we designed multiple sgRNAs targeting the gene *ADE2* with varying predicted strain coverage to create *ade2* knockout mutants in the Sardi *et al*. strain set (**Figure 3**). This strain set was chosen because we had access to the strains, but the 1011-strain population genomic data dataset (Peter *et al*. 2018) was also searched to compare relative strain coverage predictions for selected sgRNAs. Specifically, we selected two sgRNAs predicted to target all 167 strains (high-coverage sgRNAs) and two sgRNAs predicted to target only 42 of the 167 strains (low-coverage sgRNAs). The two high-coverage sgRNAs, H1 and H2, had activity scores of 1.341 and 0.426, respectively. The two low-coverage sgRNAs, L1 and L2, had activity scores of 2.569 and 2.050, respectively. None of the sgRNAs selected had any offsite matches or human hits. To determine whether the high-coverage guides also had high strain coverage within the previously published 1011-strain population genomic dataset we reran the search with the same criteria on this dataset. H1 and H2 were also predicted to cut the vast majority of the 1011-strain set, targeting 905 and 910 genomes, respectively.

**Figure 3.**
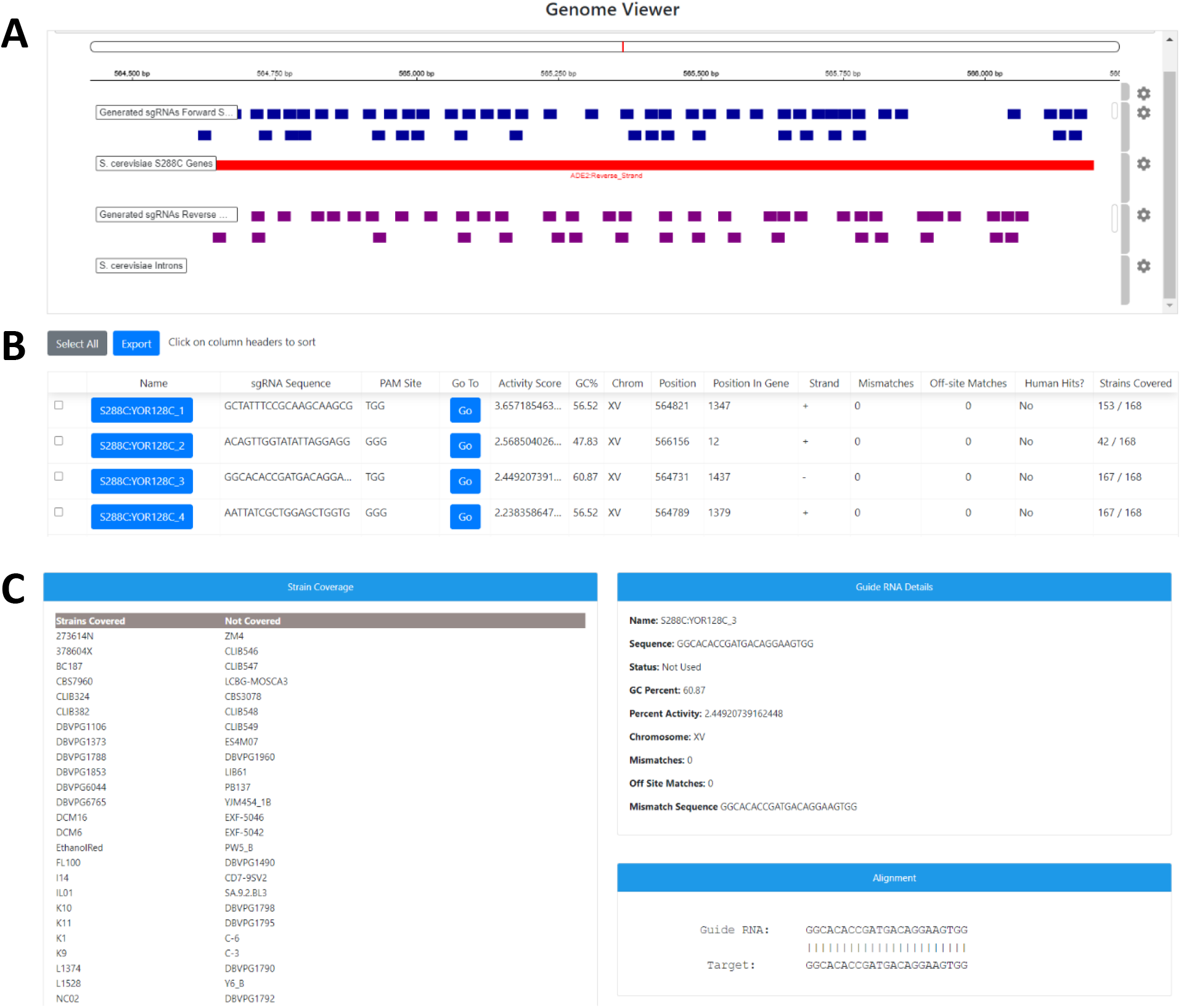
Sample output from CRISpy-pop searched for the gene *ADE2* in S288C genome with NGG PAM sequence, spacer length of 20, and cross-referencing the human genome to ensure no perfect matches exist for selected sgRNAs. A, genome viewer output by CRISpy-pop, showing the relative position of each sgRNA within the target gene. B, portion of the sgRNA table of results with each data point for each output sgRNA sequence. C, detailed results for an individual sgRNA, including identities of targeted and non-targeted strains.

### Yeast strain selection and transformations

We examined the strain coverage summary details from the CRISpy-pop search output for each sgRNA (**Figure 3C**) and selected six strains to test its predictive performance (**Figure 4**). Two strains (K1, S288C) were selected because they were predicted to be targeted by all four sgRNAs. These positive controls verified the functionality of all four sgRNAs and donor DNA constructs. The other four selected strains (L1374, SK1, T73, Y55) were predicted to be targeted by the high-coverage sgRNAs (H1 and H2) but not by the low-coverage sgRNAs (L1 and L2). S288C is haploid, while the other five strains are diploid.

**Figure 4.**
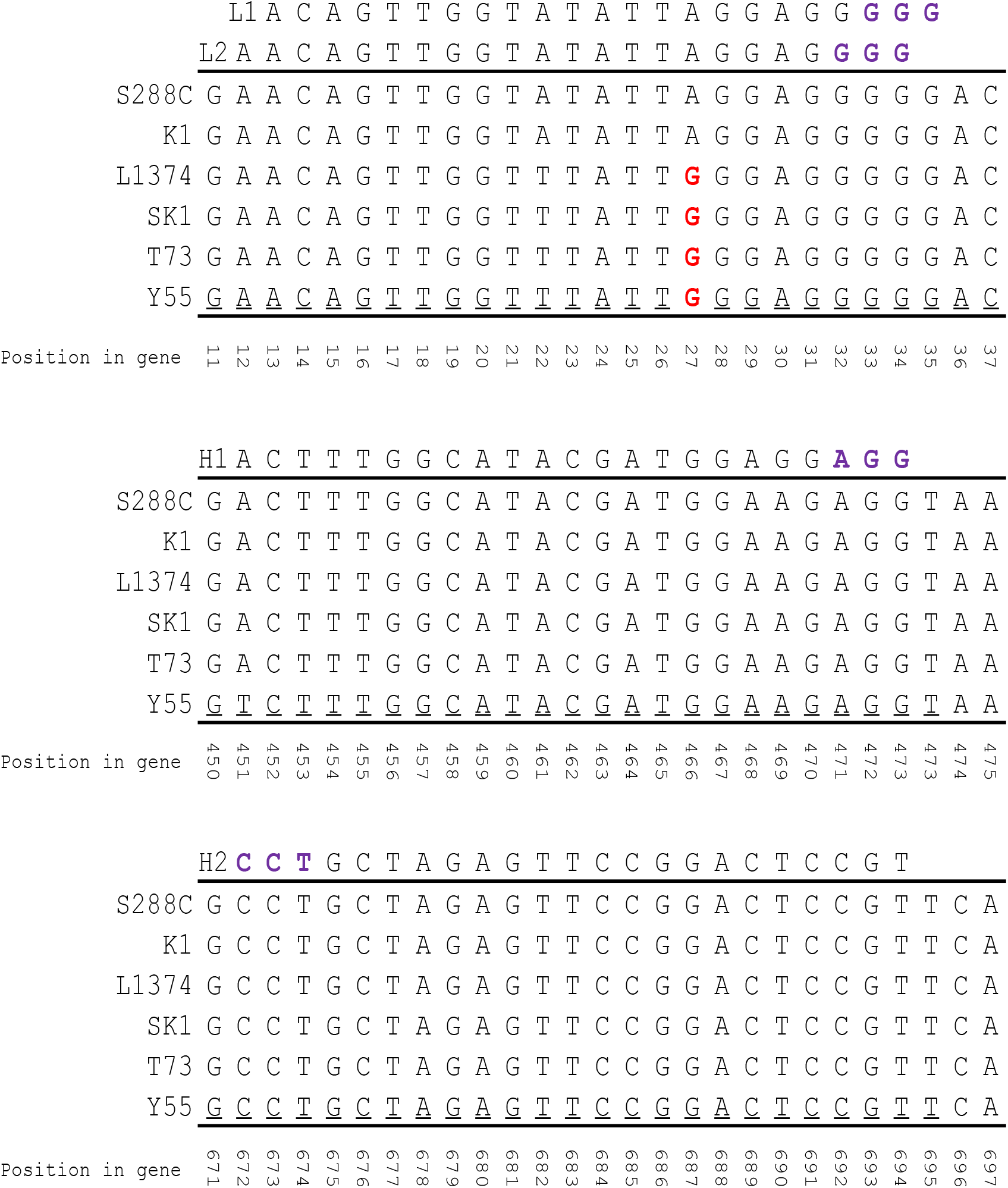
Portions of the *ADE2* gene from each strain aligned with the four sgRNAs. The PAM sites are included in purple. The *ADE2* gene sequence from each strain was extracted and aligned to each other and the four sgRNA sequences (H1, H2, L1, L2). The single nucleotide polymorphism highlighted in red at position 27 is predicted to prevent the two low-coverage sgRNAs (L1 and L2) from targeting *ADE2*. Note that sgRNA H2 targets the opposite strand, so its reverse complement is shown in this figure.

### Validation of sgRNA predictions made using CRISpy-pop

We transformed all six strains with CRISPR/Cas9 vectors expressing all four sgRNAs. We then counted the number of pink colonies, which are putative *ade2* knockouts due to deletion of *ADE2* causing the accumulation of aminoimidazole ribonucleotide (Silver and Eaton 1969), and we divided that number by the total number of transformants (G418-resistant) to calculate efficiencies (**Figure 5**). The strains that were predicted to be targeted by all four sgRNAs were transformed first to ensure that all four sgRNAs were capable of producing *ade2* knockouts. All four sgRNAs successfully targeted the two predicted strains (K1 and S288C). We verified that homology-directed repair using the donor DNA - and not NHEJ - had occurred by Sanger-sequencing the *ADE2* PCR product. Once it was confirmed that all four sgRNAs could produce *ade2* knockouts using the donor DNA, the remaining strains were transformed with all four sgRNAs and donor DNA. As predicted, the low-coverage sgRNAs did not target the four strains (L1374, SK1, T73, and Y55) predicted to only be cut by the high-coverage sgRNAs, but the high-coverage sgRNAs all resulted in *ade2* knockout mutants.

**Figure 5.**
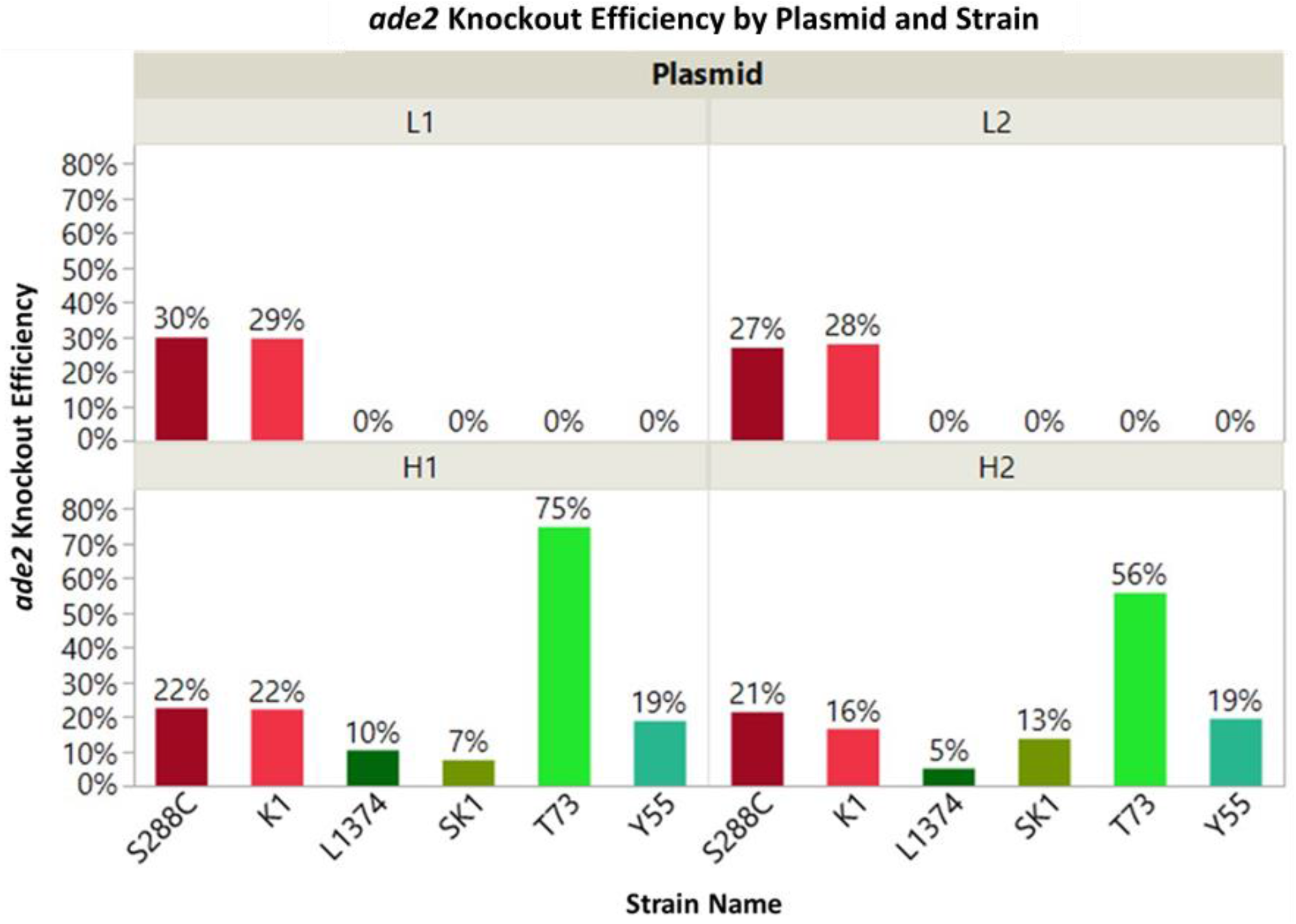
CRISpy-pop generated sgRNAs that target *ADE2* in a strain-specific manner. Results of the transformation of each strain with each sgRNA is shown. The two strains in red (S288C and SK1) were each predicted to be targeted by all four sgRNAs. Only these two strains both had non-zero %*ade2* knockouts (KOs) using all four sgRNAs. The four remaining strains were predicted to be targeted by only the high-coverage sgRNAs (H1 and H2), but not the low-coverage sgRNAs (L1 and L2). These four strains only had non-zero %*ade2* knockouts using the high-coverage sgRNAs. These results align with the strain coverage predictions made by CRISpy-pop. The predicted activity scores were H2<H1<L2<L1, which are also consistent with the observed efficiencies.

Efficiencies varied widely by strain, likely due to differences in ploidy and the relative activities of homology-directed repair. For the strains able to be cut by all four sgRNAs (K1 and S288C), the sgRNA activity scores predicted by CRISpy-pop correlated with their relative efficiencies (H2<H1<L2<L1, Kendall’s Tau = 0.85, *p* = 0.00101). These results validate the accuracy of the strain coverage and activity score predictions made by CRISpy-pop.

### Conclusions

In summary, CRISpy-pop is a powerful and flexible design tool for planning and executing CRISPR/Cas9-driven genetic modifications on individual strains or large panels of strains. CRISpy-pop can continue be expanded to support new genomes as more data become available. The ability to target different PAM sites allows potential to use or screen for other Cas systems. It correctly predicts which strains can be targeted by which sgRNAs, as well as the activities of sgRNAs. Offsite targets, including a biosafety feature that scans for potential human genome binding, can be easily avoided with CRISpy-pop. This unique combination of features and its user-friendly web interface make CRISpy-pop ideal for designing experiments in diverse populations used for genetic engineering.

## Acknowledgments

We thank Kevin S. Myers for beta testing and feedback, Maria Sardi for providing datasets, and the University of Wisconsin Biotechnology Center DNA Sequencing Facility for providing Sanger-sequencing facilities and services. This material is based upon work supported in part by the Great Lakes Bioenergy Research Center, U.S. Department of Energy, Office of Science, Office of Biological and Environmental Research under Award Number DE-SC0018409; the National Science Foundation under Grant No. DEB-1442148; USDA National Institute of Food and Agriculture (Hatch Project 1020204). C.T.H. is a Pew Scholar in the Biomedical Sciences and a H. I. Romnes Faculty Fellow, supported by the Pew Charitable Trusts and Office of the Vice Chancellor for Research and Graduate Education with funding from the Wisconsin Alumni Research Foundation, respectively.

